# Structural remodeling of the endoplasmic reticulum in response to extracellular signals requires αTAT1-induced microtubule acetylation

**DOI:** 10.1101/2023.04.20.537623

**Authors:** Hannah R. Ortiz, Paola Cruz Flores, Aaron Ramonett, Tasmia Ahmed, Nathan A. Ellis, Paul R. Langlais, Karthikeyan Mythreye, Nam Y. Lee

## Abstract

Dynamic changes in the endoplasmic reticulum (ER) morphology are central to maintaining cellular homeostasis. Microtubules (MT) facilitate the continuous remodeling of the ER network into sheets and tubules by coordinating with many ER-shaping protein complexes, although how this process is controlled by extracellular signals remains unknown. Here we report that TAK1, a kinase responsive to numerous growth factors and cytokines including TGF-β and TNF-α, triggers ER tubulation by activating αTAT1, an MT-acetylating enzyme that enhances ER-sliding. We show that this TAK1/αTAT-dependent ER remodeling promotes cell survival by actively downregulating BOK, an ER membrane-associated proapoptotic effector. While BOK is normally protected from degradation when complexed with IP3R, it is rapidly degraded upon their dissociation during the ER sheets-to-tubules conversion. These findings demonstrate a distinct mechanism of ligand-induced ER remodeling and suggest that the TAK1/αTAT pathway may be a key target in ER stress and dysfunction.

## INTRODUCTION

Endoplasmic reticulum (ER) represents the largest organelle in the mammalian cell system that extends from the nuclear envelope to the cell periphery in a vast network of tubules and sheet-like membrane domains^1–3^. It performs many diverse and highly specialized functions in part by physically interacting with other organelles including the mitochondria, lysosomes, and the nucleus as well as through active structural reorganization^4–9^. The ER sheets are the primary location for synthesis, modifications and folding of integral membrane and secreted proteins. The peripheral ER tubules are generally believed to be the main regions of lipid synthesis and signaling between the ER and other organelles. While the overall ER morphology can vary among different cell types, it is clear that ER tubules are particularly dynamic and continually rearrange to establish interconnected membranous structures throughout the cytoplasm to maintain homeostasis^10^.

Many factors contribute to ER remodeling including different classes of membrane-bending reticulons and atlastin GTPases that help fuse the membranes^1,11–13^. Equally important are cytoskeletal and motor proteins that help maintain the dynamic architecture of the ER^14,15^. Previous studies have shown that the physical association between the ER and microtubules (MTs) plays a crucial role in ER tubulation and extension along MTs, a process that often requires retrograde and anterograde machinery^14,16,17^. Two major types of MT-based ER remodeling have been described^15,18,19^. The first involves a protein complex called TAC (tip attachment complex) in which the tip of an ER tubule is linked to the plus end of an MT and the ER tubule extends or regresses together^19^. The second type is ER sliding wherein a new tubule is pulled from existing tubules along the length of a stable MT. Studies have shown that ER sliding is the most frequently observed type of ER movement and preferentially occurs along acetylated MTs though the underlying reasons remain unclear^16,19,20^.

However, given that ER dynamics occur mostly along a subset of acetylated MTs, and that the overall function of ER sliding is poorly defined, we were motivated to explore how TAK1, a serine/threonine kinase activated by members of the transforming growth factor β (TGF-β) superfamily as well as inflammatory cytokines and stress conditions, influences ER remodeling and functions. Indeed, we previously reported that TAK1 directly regulates α-tubulin acetyltransferase 1 (αTAT1) activity, the sole enzyme responsible for MT acetylation *in vivo*^21–28^. Notably, we showed that TAK1 phosphorylates αTAT1 at Ser237, which activates this enzyme to dramatically enhance MT acetylation in most cell types^21^.

In the present study, we report that TGF-β signaling can cause rapid ER remodeling through the TAK1/αTAT1/MT acetylation pathway. Moreover, we demonstrate that a distinct role of ER tubulation in regulating cell survival through turnover of a key proapoptotic effector.

## RESULTS

### TAK1 activation causes ER remodeling into peripheral tubules

Based on our previous discovery that TAK1 directly activates αTAT1^21^, we hypothesized that this pathway may in turn promote MT acetylation-dependent ER remodeling. We began exploring this possibility by first monitoring for dynamic changes in ER morphology in response to various TAK1 inducers such as TGF-β and TNF-α. COS-7 cells were briefly stimulated with TGF-β prior to fixation at different time points and dual stained for ER markers: reticulon-4 (NOGO; green), an ER-sculpting protein distributed throughout the ER, and calnexin (red), which preferentially localizes in the lumen of the perinuclear rough ER (Fig. 1A, Fig. S1A). Immunofluorescence analysis revealed a rapid shift in ER morphology from largely sheet-like appearance to more tubulated networks, as indicated by NOGO staining, within 5 min and plateaued upon 30 min (Fig. 1A, graph). Similar effects were observed in another cell line, Panc1, wherein TGF-β and TNF-α were each capable of enhancing ER tubulation within minutes (Fig. 1B, upper panels; Fig. S1B). To determine whether these effects were specifically TAK1-dependent, a parallel study was performed in which TAK1 expression was stably knocked down in Panc1s via shRNA-mediated gene silencing (Fig. S1C). Here, TGF-β stimulation again induced peripheral ER tubulation in control Panc1 but not in TAK1 knockdown cells (shTAK1), which instead, displayed more densely packed ER sheets along the perinuclear and peripheral space irrespective of TGF-β stimulation (Fig. 1B, lower panels and graph). Inhibition of the TAK1 kinase with a small molecule also yielded a diffuse, sheet-like ER morphology independent of TGF-β treatment (Fig.1C), thus indicating that ER remodeling induced by TGF-β and TNF-α requires TAK1 activity.

**Fig. 1.**
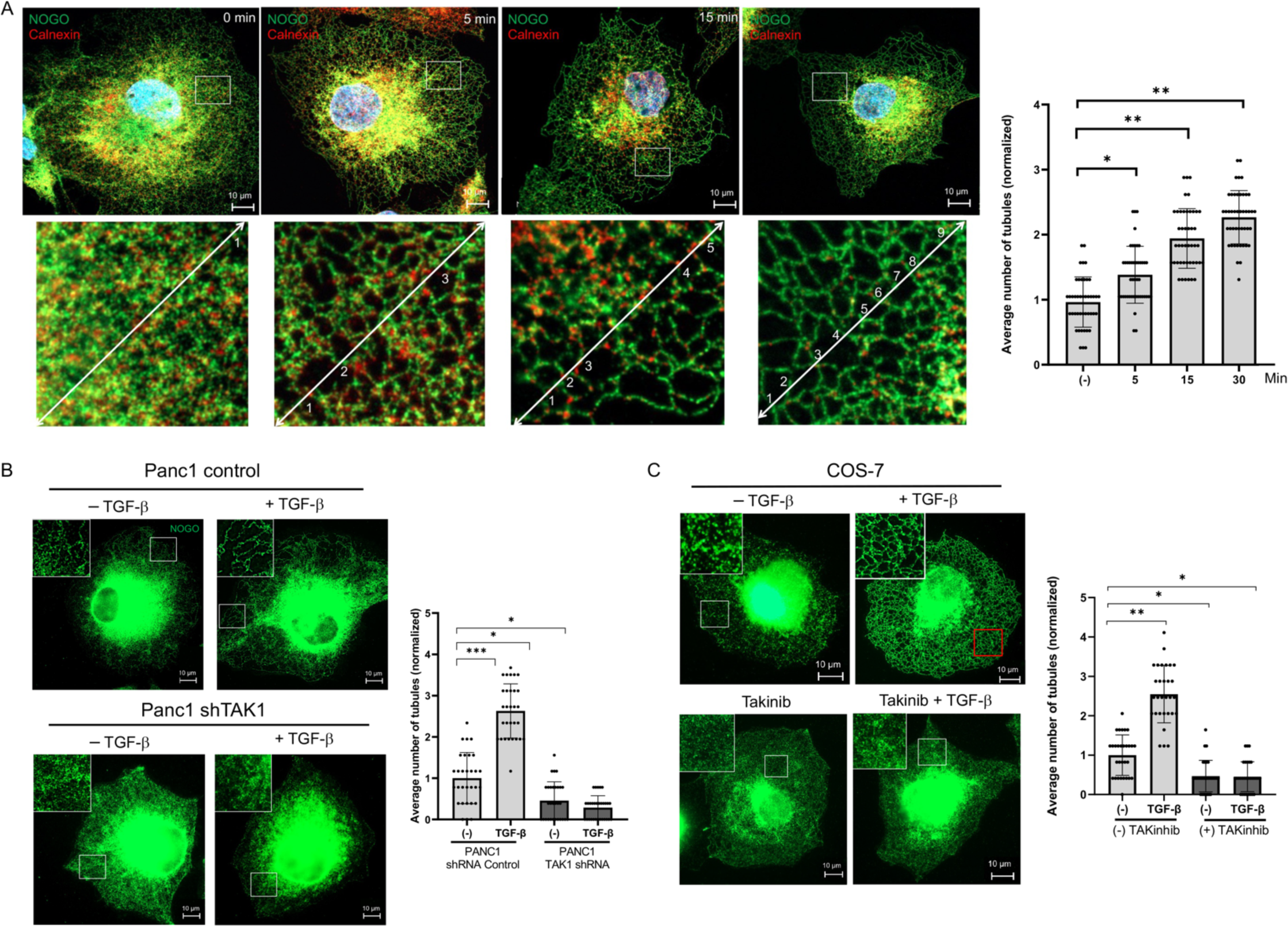
TAK1 activation causes ER remodeling into peripheral tubules. A) COS-7 were treated with TGF-β (200 pM) for up to 30 min, fixed and imaged for reticulon 4 (NOGO) and calnexin immunofluorescence staining. Graph depicts the increase in ER tubulation upon TGF-β induction. ER tubulation was quantified using FIJI plot profile using 3 random ROIs per cell (20 cells) and counting peaks showing greater differences in amplitude. B) Representative images for immunofluorescence staining of NOGO in control Panc1 and Panc1 shTAK1 cell line treated with and without TGF-β (200 pM) for 30 min, then fixed and imaged for NOGO immunofluorescence staining. Graph represents ER tubulation. C) COS-7 cells were treated with and without TGF-β (200 pM) for 30 min and with and without pretreatment of TAKinib (10 uM) for 30 min then fixed and imaged for Reticulon-4 immunofluorescence staining. *p<0.05. Error bars represent SEM. *p<5×10^−4^ and **p< 5×10^−12^ relative to control.

### ER remodeling requires αTAT1-induced acetylation and stabilization of MTs

Although there exists a clear link between MT acetylation and ER sliding^16^, how these two processes are coordinated by extracellular cues remains unclear. Based on our observation of TAK1-mediated ER remodeling, we next tested whether these effects require MT acetylation. Consistent with our previous findings^21^, TGF-β stimulation greatly enhanced MT acetylation within minutes (red), which coincided with increased ER tubulation (green) (Fig. S2). Importantly, a significant overlap was observed for ER tubules along acetylated MTs upon TGF treatment (second panel, yellow colocalization indicated by white arrows), whereas pharmacologic inhibition of the TAK1 kinase abrogated these responses (Fig. S2; lower two panels), suggesting that the TGF-β/TAK1-induced MT acetylation critically promotes ER sliding and tubulation. Therefore, to test whether αTAT1 is a key mediator of these TAK1-dependent effects, we utilized αTAT1 knockout (αTAT1-KO) mouse embryonic fibroblasts (MEF). Here, TGF-β again triggered a robust response in peripheral ER tubulation in control MEFs whereas αTAT1-KO maintained a mostly sheet-like distribution irrespective of ligand treatment (Fig. 2A; graph).

**Fig 2.**
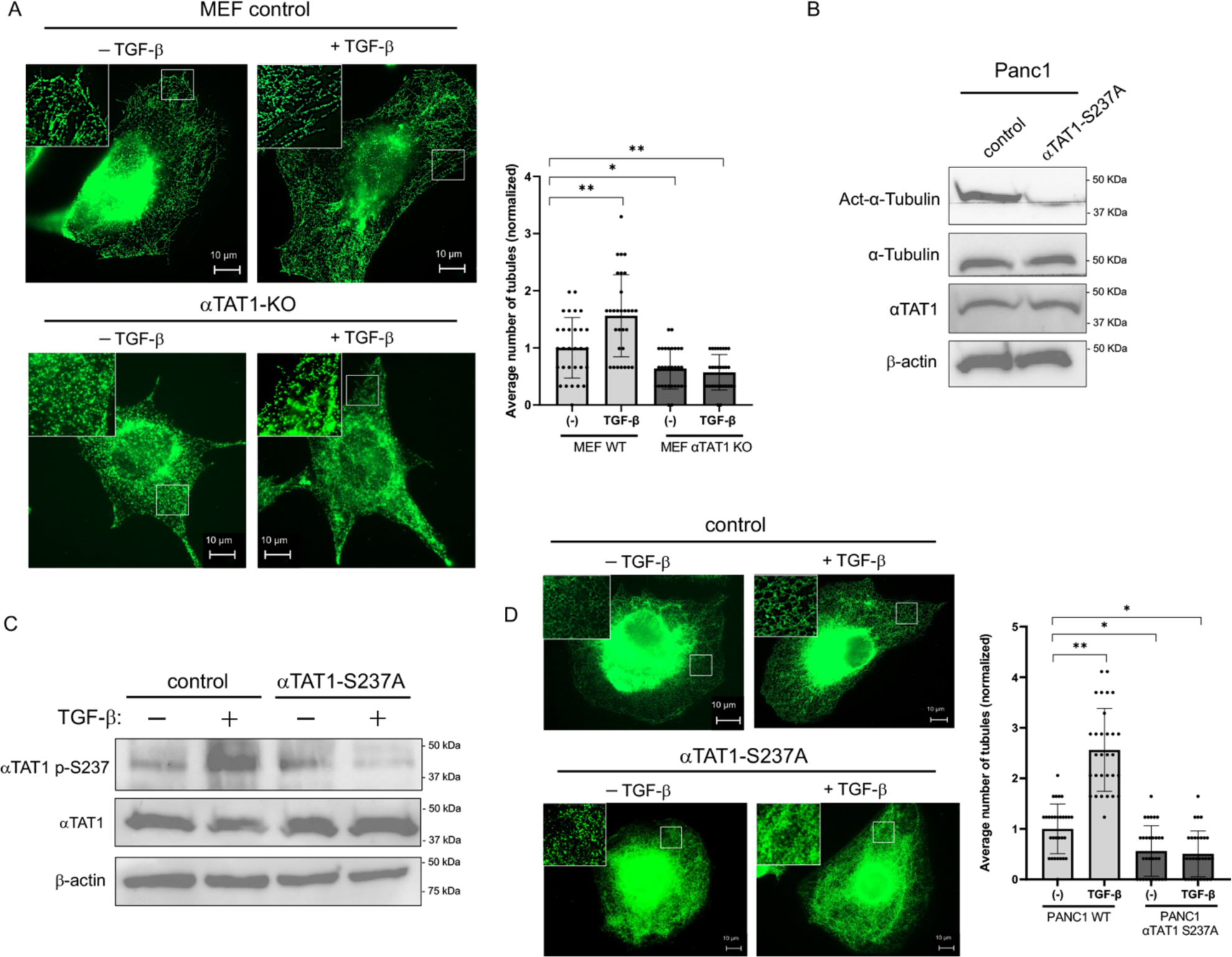
ER remodeling requires αTAT1-induced acetylation and stabilization of MT. A) Representative images of immunofluorescence staining of NOGO in MEF WT and αTAT1 KO cells, treated with and without TGF-β (200 pM) for 30 min prior to staining. *p< 5×10^−2^ and **p< 5×10^−3^ relative to control. B) Western blots for Panc1 WT and αTAT1 S237A cells show endogenous expression of acetyl-α-tubulin, α-tubulin, αTAT1 and β-actin levels. C) Western blots for Panc1 WT and αTAT1 S237A cells subjected to 30 min TGF-β (200 pM) showing αTAT1 S237 phosphorylation using a phosphoantibody as well as levels of total αTAT1 and β-actin. D) Representative images of immunofluorescence staining of NOGO in Panc1 WT and Panc1 αTAT1 S237A cells, treated with and without TGF-β (200 pM) for 30 min prior to staining. Graph represents ER tubulation quantified using FIJI plot profile using 3 respective ROIs for 20 cells and counting peaks showing greater differences in amplitude, *p< 5×10^−2^ and **p< 5×10^−24^ relative to the control.

The above results prompted us to explore how defects in TAK1-induced αTAT1 activation specifically alters ER dynamics. To do so, a knock-in cell line was generated harboring a single amino acid substitution in αTAT1 that renders the enzyme nearly inactive. Indeed, we previously identified Ser237 of αTAT1 as a powerful regulatory site, which upon phosphorylation by TAK1, dramatically enhances MT acetylation by up to 80-90% over basal levels in various cell types^21^. Accordingly, three clones were generated in Panc1 cells through CRISPR/Cas9 genomic editing, each of which carried the endogenous alanine knock-in mutation (αTAT1-S237A) but expressing similar levels as wildtype (WT) control cells and displaying defects in MT acetylation as shown by biochemical and immunofluorescence analyses of α-tubulin acetylation (Fig.2B, S3A-C). Based on these characterizations, we next tested whether this αTAT1 phosphorylation site is critical for ER-tubulation. We began by utilizing our αTAT1-S237 phosphoantibody, which we previously determined to be compatible with biochemical assessment of αTAT1 activity and found that TGF-β strongly induces αTAT1 activation in control Panc1 cells but not in S237A knock-in or shTAK1 cells as expected (Fig.2C and S4A). More importantly, immunofluorescence studies revealed a decisive role for TAK1 in ER remodeling through αTAT1 activation as both TGF-β and TNF-α induced peripheral ER tubulation in control but not S237A cells (Fig. 2D and S4B).

Because of recent evidence of MT acetylation conferring mechanical stability against breakage and ageing^27,29–31^, we next tested whether the αTAT1-mediated ER tubulation is primarily due to the increased stability of MTs. A pharmacological approach was used starting with paclitaxel, a MT stabilizing agent, which markedly enhanced ER tubulation in Panc1 control cells independent of TGF-β treatment whereas it had little effect on the ER morphology of αTAT1-S237A cells (Fig. 3A; graph). Conversely, we chemically disrupted MT stabilization using nocodazole to find that ER remodeling was profoundly impaired in both WT and αTAT1-S237A cells independent of ligand stimulation (Fig. 3B; graph). Collectively, these results indicated that both the acetylation and structural stability are important requirements of ER remodeling in response to extracellular stimuli.

**Fig 3.**
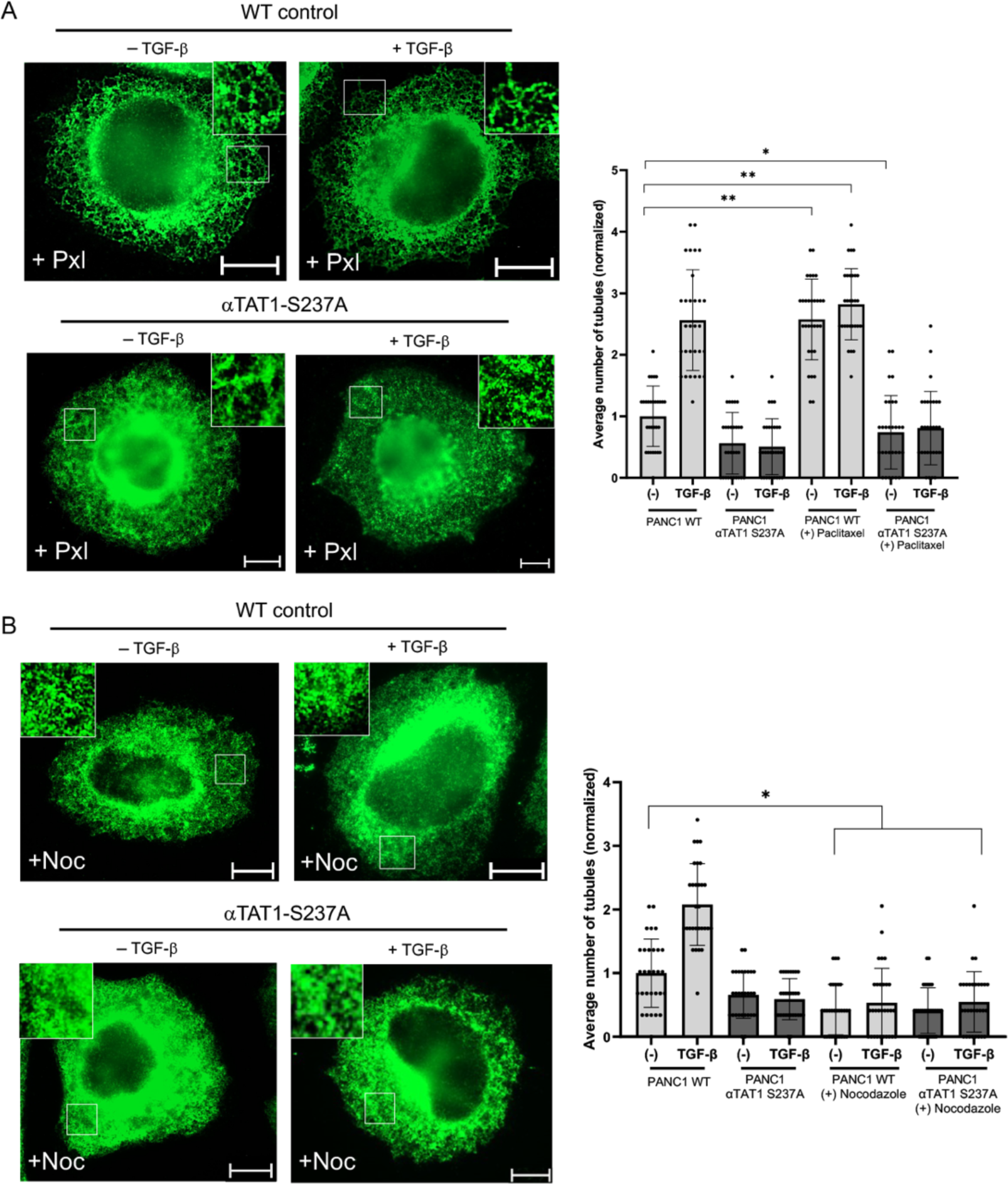
TGF-β driven TAK1 αTAT1 phosphorylation and MT stability drives endoplasmic reticulum remodeling. A) Representative images of immunofluorescence staining of NOGO in Panc1 WT and αTAT1 S237A cells, pretreated with paclitaxel (100 nM) for 1 hour prior to TGF-β stimulation (200 pM) for 30 min before staining. Graph represents ER tubulation quantified using FIJI plot profile using 3 respective ROIs for 20 cells and counting peaks showing greater differences in amplitude. *p<0.05 and **p< 5×10^−11^ relative to the control. B) Representative images of immunofluorescence staining of NOGO in Panc1 WT and αTAT1 S237A cells, pretreated with nocodazole treatment (100 nM) prior to TGF-β stimulation (200 pM) for 30 min prior before staining. Graph represents ER tubulation quantified using FIJI plot profile using 3 respective ROIs for 20 cells and counting peaks showing greater differences in amplitude. *p< 5×10^−11^ relative to the control.

### TAK1/αTAT1-mediated ER remodeling regulates apoptosis through BOK turnover

To identify functional roles associated with the TAK1/αTAT1-dependent ER remodeling, we performed mass spectrometry-based quantitative proteomics on control and αTAT1-S237A cells (Fig. 4A and S5). Data showed the differential regulation of 224 proteins, among which 130 were associated with the ER based on gene ontology analysis (Fig. 4B, C). Upon further statistical analysis, we noted BCL2-related ovarian killer (BOK) as a lead candidate as it was one of the most highly upregulated proteins in αTAT1-S237A cells (Fig. S5). Subsequent biochemical analysis confirmed our proteomics data, as shown by at least a three-fold increase in BOK level in the mutant cell line compared to control (Fig. 4D). Consistent with the biochemical results, immunofluorescence analysis revealed much greater levels of BOK (red) at basal state in αTAT1-S237A cells than WT control (Fig. 4E; graph). Interestingly, there was a notable reduction in BOK levels upon TGF-β stimulation in control but not αTAT1-S237A cells (Fig. 4E), suggesting that BOK expression is regulated by ER remodeling.

**Fig 4.**
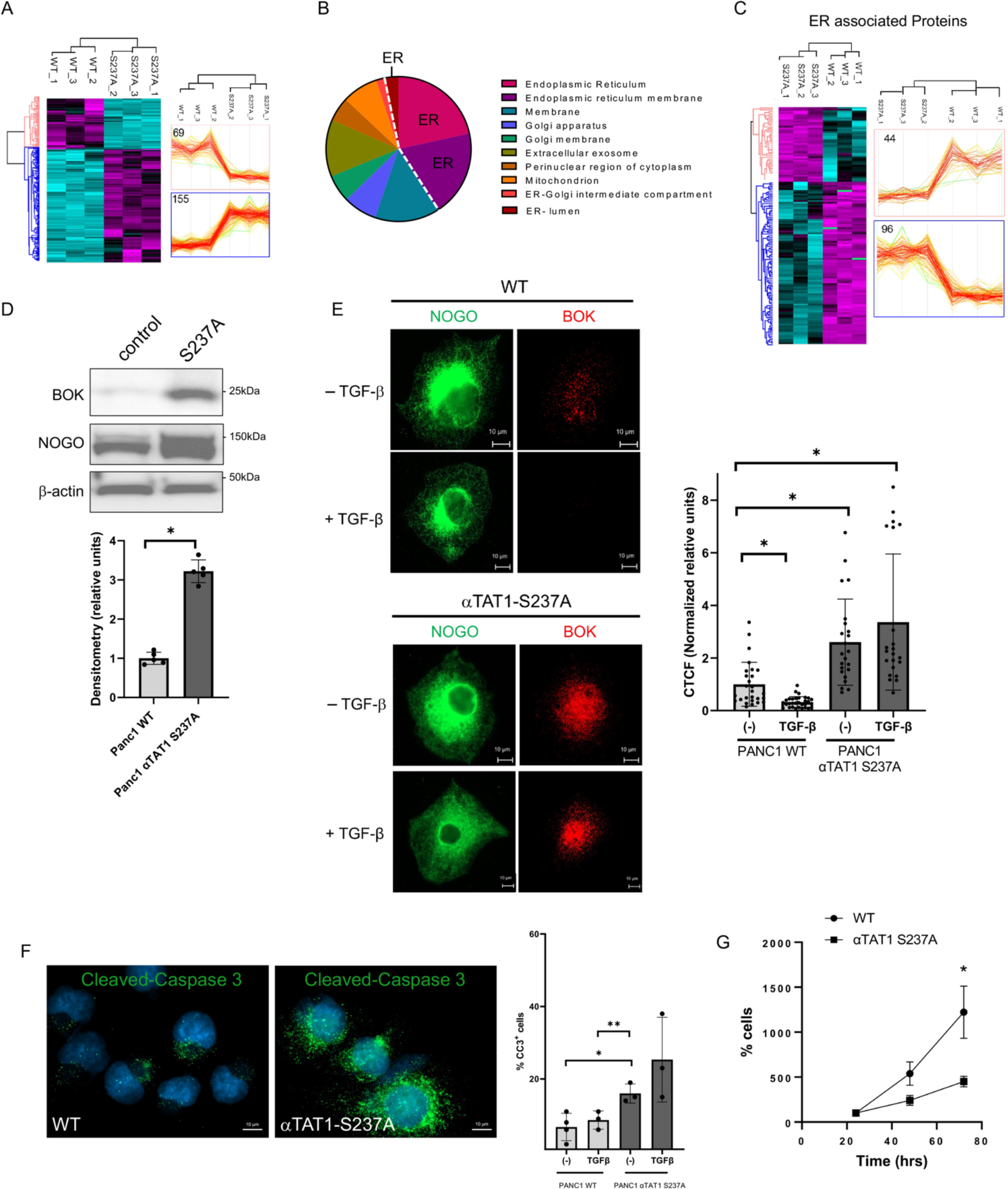
TAK1/aTAT1-mediated ER remodeling regulates apoptosis through BOK turnover. A) Heat map based on MS-based quantitative proteomics shows differential expression of proteins found in Panc1 WT versus αTAT1 S237A. N=3 biological replicates per group. B) Pie chart depicts the cellular comparts from gene ontology data from MS-based quantitative proteomics. C) Heat map based on MS-based quantitative proteomics shows differential expression of ER associated proteins found in Panc1 WT versus αTAT1 S237A. D) Western blots for Panc1 WT and αTAT1 S237A cells show endogenous expression of BOK, NOGO and β-actin levels, where BOK levels measured from n= 5 independent experiments. Error bars represent SEM, *p<5×10^−5^. E) Representative images of immunofluorescence staining of NOGO (Green) and BOK (Red) in Panc1 WT and αTAT1 S237A cells treated with and without TGF-β (200 pM) for 30 min prior to staining. Graph represents CTCF quantification of BOK taken from an average of three independent experiments where at least 22 cells were quantified per presented as a mean value ± SEM. A Type 2 t-test results show: *p<0.0005. F) Representative images of immunofluorescence staining of cleaved-caspase-3 in Panc1 WT and αTAT1 S237A cells. Graph represents the positive number of cleaved-caspase-3 cells taken from an average of three independent experiments where at least 20 cells were quantified per group per experiment presented as a mean value ± SEM. A Type 2 t-test results show: *p=0.05. G) Crystal violet assay shows comparison of proliferation between Panc1 WT and αTAT1 S237A cells over 72 h. Graph represents normalized average from three independent experiments. Error bars represent SEM and *p=0.05.

Since BOK is a proapoptotic BCL2 protein^32–37^, we tested whether the aberrant rise in BOK expression in αTAT1-S237A cells promotes apoptosis by monitoring the cleaved caspase-3 levels. As expected, there was a two to three-fold increase in the percentage of cells displaying caspase cleavage (green) in the mutant cells relative to control at basal state although TGF-β appeared to have little effect on either group at least during the 2 h treatment (Fig. 4F; graph). However, consistent with the enhanced caspase cleavage, αTAT1-S237A cells displayed a markedly impaired growth rate relative to control (Fig. 4G), suggesting that the αTAT1 regulation of BOK expression is a key determinant of cell survival.

### Dynamic BOK turnover requires TAK1-induced αTAT1 phosphorylation

Based on the observation that BOK levels rapidly decrease following TGF-β stimulation, we surmised that this loss is primarily due to enhanced protein turnover. We conducted biochemical assays to test this and further explore how the TGF-β-induced TAK1/αTAT1 activation is linked to BOK degradation. First, we compared BOK expression in control and TAK1 knockdown stable cells upon a brief TGF-β stimulation to find a concentration-dependent BOK attenuation in control cells which inversely correlated with enhanced αTAT1 phosphorylation (Fig. 5A; top and middle panels and graph). In contrast, the observed lack of αTAT1 activation in shTAK1 cells corresponded strongly with BOK upregulation at basal state and post-ligand treatment (Fig. 5A), suggesting that TAK1-dependent αTAT1 activation is necessary for efficient BOK turnover. To further prove that TAK1 phosphorylation of αTAT1 at S237 is specifically required, we conducted a parallel experiment using αTAT1-S237A cells to again find that BOK levels decrease with increasing TGF-β concentrations in control cells in an αTAT1-dependent manner whereas S237A mutant cells maintained high levels of BOK independent of TGF-β treatment (Fig. 5B; graph).

**Fig 5.**
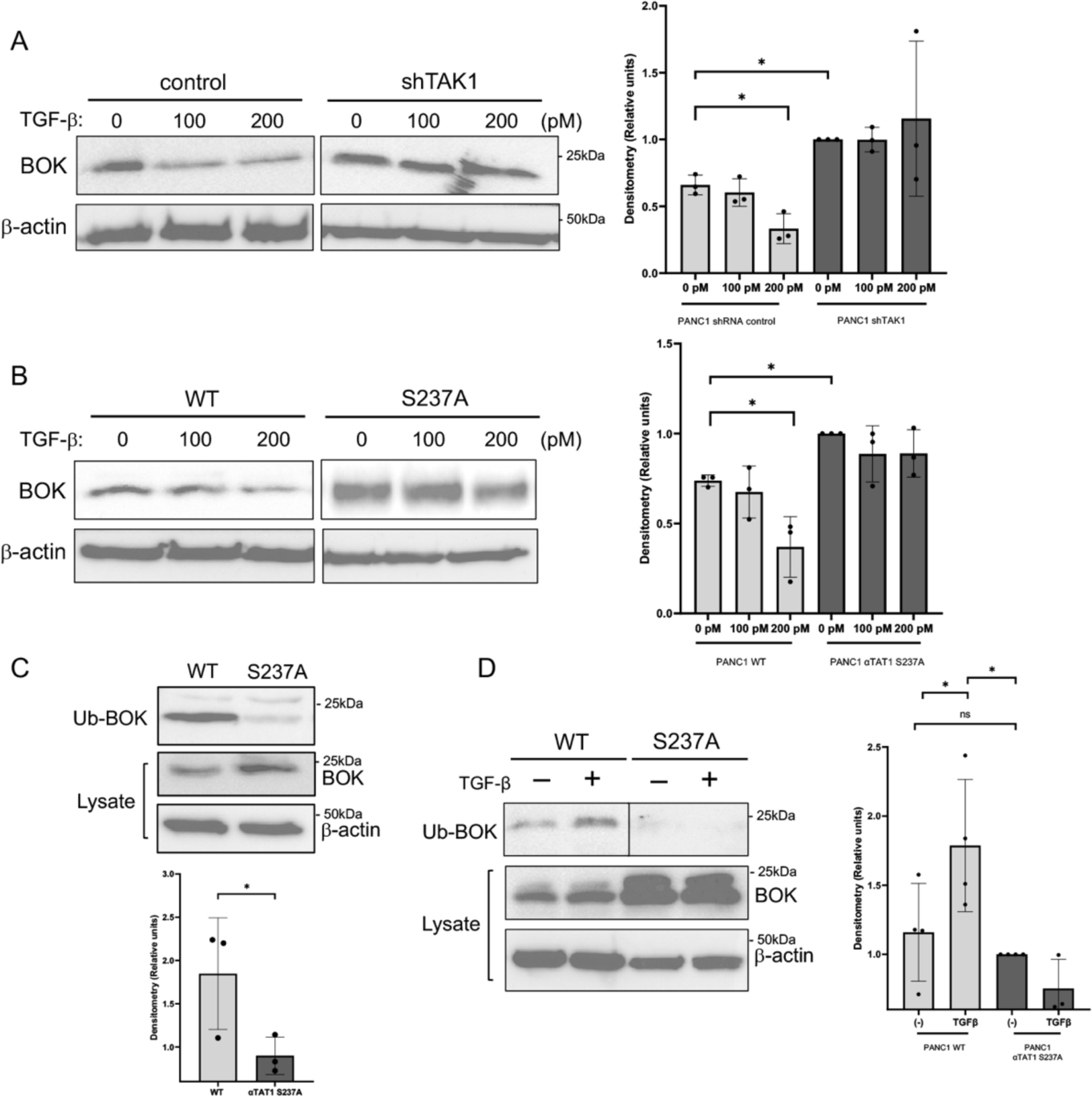
TAK1-induced αTAT1 phosphorylation regulates BOK turnover. A) Panc1 WT and shTAK1 cell were treated with increasing concentrations of TGF-β (0, 100 and 200 pM) for 30 min before western blot analysis of BOK and β-actin. Graph represents quantification for BOK levels as an average of three independent experiments. Error bars represent SEM and *p=0.05. B) Panc1 WT and αTAT1 S237A cell were treated with increasing concentrations of TGF-β (0, 100 and 200 pM) for 30 min before western blot analysis of BOK and β-actin. Graph represents quantification for BOK levels as an average of three independent experiments. Error bars represent SEM and *p=0.05. C) Panc1 WT and αTAT1 S237A were immunoprecipitated with ubiquitin antibody. Graph represents quantification of BOK levels from three independent experiments. Error bars represent SEM and *p=0.05. D) Panc1 WT and αTAT1 S237A cells were treated with TGF-β (200pM) for 1 hr before immunoprecipitation with ubiquitin antibody. Graph represents quantification of BOK levels from three independent experiments. Error bars represent SEM and *p=0.05.

As BOK have been previously shown to be degraded via the ubiquitin proteosome pathway^34^, we hypothesized that there would be reduced BOK ubiquitination in S237A mutant cells relative to control. Indeed, results showed a dramatic reduction in ubiquitinated BOK in αTAT1-S237A cells despite having a much greater total level of BOK (Fig. 5C). Moreover, BOK ubiquitination increased upon TGF-β treatment in control cells whereas the S237 cells remained unresponsive (Fig. 5D), further supporting the notion that BOK turnover via the ubiquitin proteosome pathway is critically dependent on αTAT1 phosphorylation by TAK1.

### BOK turnover is mediated by TAK1/αTAT1-dependent disruption of BOK/IP3R interaction during ER remodeling

Lastly, we sought to determine how MT acetylation-dependent ER remodeling mediates BOK turnover by considering the reported link between BOK and the inositol 1,4,5-trisphosphate receptor (IP3R). Previous studies have shown that BOK protein is stabilized at basal state when physically bound to IP3R on the ER membrane surface, whereas unbound BOK gets degraded via the ubiquitin proteosome pathway^36,37^. Accordingly, we hypothesized that the TGF-β-induced BOK degradation is caused by the uncoupling of the BOK/IP3R complex during ER remodeling. In testing this mechanism, we first observed that, at basal state, BOK mostly localizes to the perinuclear ER regions and the nucleus but is absent along the peripheral ER tubules in control cells (Fig. 6A; BOK in red, NOGO in green, BOK/NOGO colocalization in yellow clusters within red boxes and further indicated by red arrows in zoomed-in gray scale). As expected, there was a significant level of BOK turnover following TGF-β stimulation although the remaining nondegraded pool remained stationary during ER tubulation (Fig. 6A; indicated by red arrow of white clusters). In S237A mutant cells, BOK accumulated in the condensed perinuclear ER domains and persisted irrespective of TGF-β treatment (Fig. 6A; graph), suggesting that the BOK stability is controlled in part by the structural remodeling of the ER (Fig. 6A). In contrast, IP3R proved highly dynamic as it translocated from the perinuclear ER to the cell periphery following TGF-β induction in control but not in S237 cells (Fig. 6B; graph). Instead, both IP3R and BOK were enriched within the dense perinuclear ER sheets in the mutant cells, hence suggesting that ER tubulation regulates the BOK/IP3R interaction and BOK protein stability. In support of this putative mechanism, BOK distribution was visualized in the presence of MG132 and found that its protein level was greatly elevated but still confined to the perinuclear ER while IP3R dispersed to the cell periphery upon TGF-β treatment (Fig. S6).

**Fig 6.**
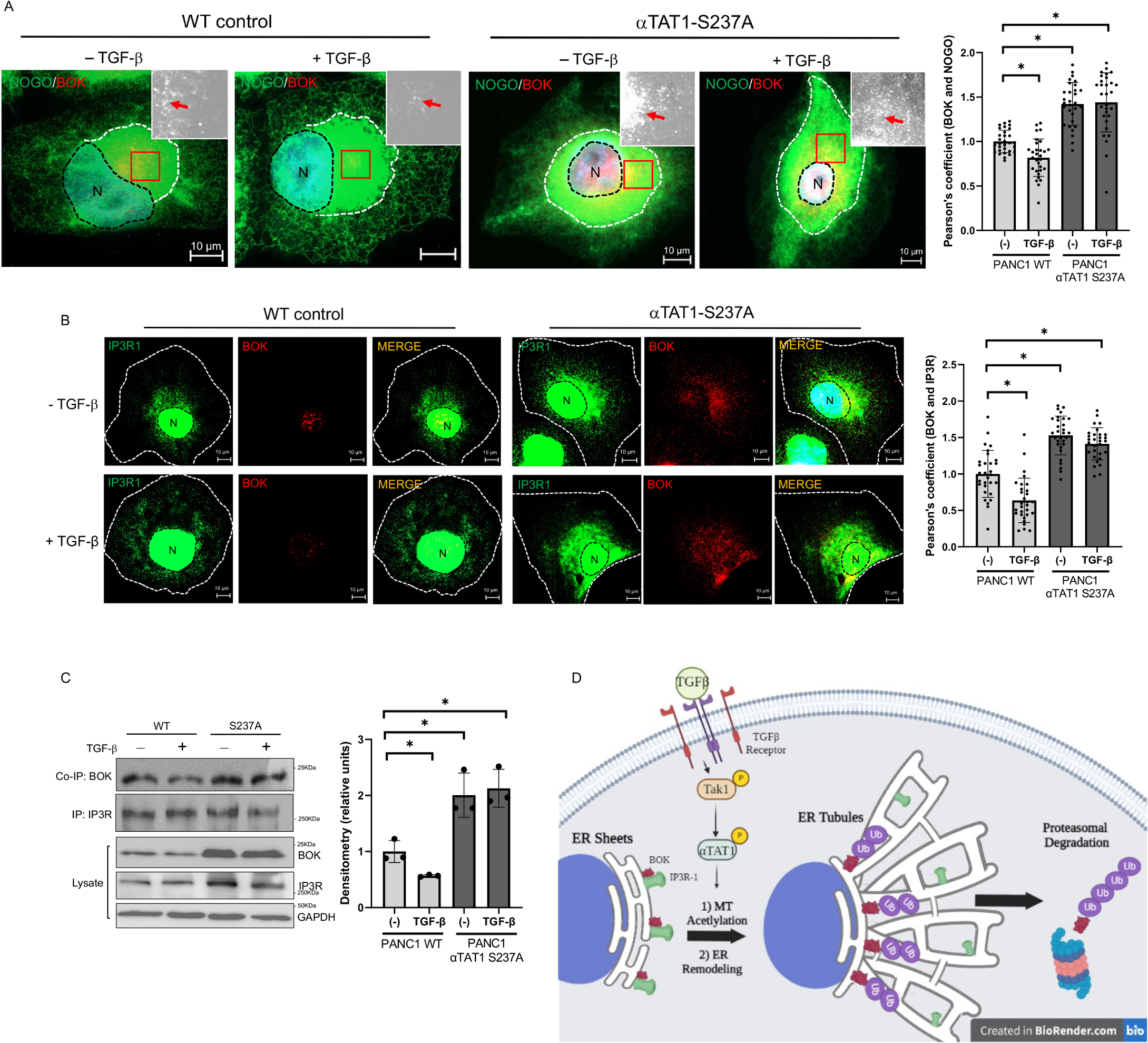
TAK1/ αTAT1-dependent disruption of BOK/IP3R interaction during ER remodeling mediates BOK turnover. A) Representative images of immunofluorescence staining of NOGO (green) and BOK (Red) in Panc1 WT and S237A cells treated with or without TGF-β (200 pM) for 30 min prior to staining and quantification based on Pearsons correlation. Colocalization of BOK/NOGO are visualized in yellow clusters and as white dots indicated by red arrows in zoomed in gray scale images. Nuclear regions are shown by a black dotted line and perinuclear endoplasmic reticulum sheets are located within the white dotted line. Graph represents the average of at least 20 cells per group with 3 ROIs per cell. *p<5×10^−5^ relative to the control. B) Representative images of immunofluorescence staining of endogenous IP3R (green) and BOK (Red) panc1 WT and αTAT1-S237A, treated with and without TGF-β (200 pM) for 30 min prior to staining. Quantification based on Pearsons correlation coefficient. Graph represents the average of at least 20 cells were quantified per group with 3 ROIs per cell. *p<5×10^−5^ relative to the control. C) Panc1 WT and αTAT1 S237A cells were treated with TGF-β (200 pM) for 1 hr before immunoprecipitation with IP3R antibody. Graph represents quantification of the BOK CO-IP levels from three independent experiments. Error bars represent SEM and *p=0.05. D) Schematic representation of BOK turnover mediated by TAK1/αTAT1-dependent disruption of BOK/IP3R interaction during ER remodeling.

To further confirm that the TGF-β-induced IP3R translocation is specifically driven by TAK1, we performed a parallel experiment in COS7 cells and found that TGF-β promotes the anterograde movement of IP3R from mostly perinuclear domains to the cell periphery (Fig. S7; top two panels) whereas TAK1 inhibition effectively blocked this anterograde redistribution (Fig. S7; lower two panels). A similar pattern was observed in Panc1 cells wherein the TGF-β-responsive IP3R translocation was abrogated in shTAK1 cells (Fig. S8; graph), thus demonstrating that TAK1 is a critical upstream regulator of the BOK/IP3R interaction and BOK stability.

Finally, to confirm that the TGF-β/TAK1 pathway regulates BOK stability by physically disrupting the interaction between BOK and IP3R during ER tubulation, we performed a co-immunoprecipitation and observed that their basal interaction was attenuated following a brief TGF-β stimulation in control whereas it was markedly higher in S237A cells at both basal state and upon TGF-β treatment (Fig. 6C; graph). Taken together, these results supported a distinctly new cellular process in which our previously reported mechanism of TAK1-induced αTAT1 activation and MT acetylation control ER tubulation and BOK stability (Fig.6D).

## DISCUSSION

Despite the growing number of ER-shaping proteins and cytoskeletal machinery that have been identified, surprisingly little is known about how ER remodeling occurs in response to various extracellular cues. The present study demonstrates what are believed to be the first identified extracellular signals that drive real-time changes in ER morphology, a process mediated by the activation of an MT-acetylation enzyme that enhances ER-sliding and tubulation.

We begin our discussion by highlighting the development of two novel reagents to investigate the role of a key αTAT1 phosphoregulatory site-the S237 phosphoantibody and a S237A knock-in cell line, which combined provided a direct read-out of αTAT1 activity and MT acetylation in response to TGF-β and TNF-α stimulation. While previous studies have relied on gene knockout, knockdown and overexpression systems to interrogate αTAT1 function^22–26,28,38,39^, this new approach represents a distinctly new method to specifically gauge for acetyltransferase versus deacetylase-dependent effects on MT acetylation (i.e. αTAT1 vs HDAC6 or SIRT2). Moreover, these reagents may have important diagnostic or prognostic implications. For instance, although some studies have linked αTAT1 to tumor growth and metastasis in various cancer cell types^22,23^, others including several tissue arrays have shown very little correlation, as its protein expression was detected at weak to moderate levels in most cancers and normal tissues^27,40–42^. Therefore, assessing for altered αTAT1 activity may offer a new approach in determining which cancer types possess αTAT1 hyperactivity and are therefore most susceptible to targeted inhibition. While beyond the scope of this study, efforts are underway to screen for S237 phosphorylation across different human cancers.

Aside from the notable reagents used in the study, one of our major findings is that the TAK1/αTAT1-dependent ER remodeling inhibits apoptosis by targeting BOK, which appears to be rapidly degraded upon its dissociation from IP3R during ER conversion from perinuclear sheets to tubules. How this physical reshaping of the organelle dictates their molecular interaction remains to be determined although it is clear that IP3R, but not BOK, can rapidly translocate from perinuclear sheets to ER tubules in response to extracellular signals. Indeed, although previous studies have indicated that BOK is protected from degradation when in complex with IP3R complex, our data explicitly shows that this interaction occurs mostly within the perinuclear ER regions at basal state in different cell types. But in response to extracellular signals, IP3R translocates to the cell periphery by tethering along with the expanding ER tubular networks. This demonstrates that the precise spatial distribution of IP3R is a crucial factor in BOK stability and turnover, and that their interaction may not be as constitutive as once thought.

The above findings then prompt new questions about how BOK actually separates from IP3R during ER tubulation. Earlier studies have defined several structural elements mediating the BOK/IP3R interaction. In particular, the BH4 helical segment found in BOK was shown to dock with a small surface exposed loop within the ARM3 domain of IP3R, a conserved and highly charged domain that contains multiple regulatory sites including for proteolysis^37^. These structural elements are currently believed to facilitate a direct and constitutive interaction between the two proteins although the potential role of other intermediates cannot be fully ruled out. It is possible that these binding elements are subjected to posttranslational modifications (PTMs) that trigger local conformational changes to promote the disruption of the BOK/IP3R interaction. In this scenario, it would be beneficial to determine how TAK1/αTAT1 activation specifically influences BOK and IP3R PTMs aside from their ubiquitination. As previous data suggest that BOK, for instance, is modified by PKA and GSK3-α/β^43^, both of which are known to crosstalk with TAK1 signaling, it would be worthwhile to explore how PTMs in BOK and IP3R are linked to TGF-β and TNF-α-dependent ER remodeling.

BOK has been considered a tumor suppressor as its downregulation has been reported in a wide variety of human cancers^34,44,45^. Consistent with these reports, our data demonstrate that TAK1/αTAT1 pathway plays a significant role in cell growth and survival by regulating BOK levels and thereby suppressing apoptosis. The fact that a single knock-in point mutation on αTAT1 results in such powerful ER-remodeling defects and subsequent increase in BOK-induced apoptosis suggest that the components of the TAK1/αTAT1 pathway may be effective therapeutic targets in certain cancers.

Finally, while we chose to focus on BOK as the top responder immediately impacted by defective ER remodeling, our quantitative proteomics data yielded numerous other responders that have yet to be investigated. Among the ER-related candidates, we note proteins such as ERAP1 and EMC9, which are linked to antigenic peptide processing and protein insertion into ER membranes, respectively. How these and other candidates are involved in the many diverse functions of this organelle including ER associated protein degradation and other stress response remain to be explored.

In summary, our present findings demonstrate a distinct mechanism by which ER remodeling occurs in response to extracellular signals. This dynamic response induced by the TAK1/αTAT1 pathway that alters MT acetylation has important implications with respect to many ER-related malignant, metabolic, and neurodegenerative diseases.

## METHODS

### Western blotting

Cell lysates were collected and separated by SDS-PAGE and through electrophoresis were transferred onto a PVDF (Polyvinylidene difluoride) membrane (BIO-RAD). The membranes were blocked with 5% skim milk in TBS with 0.1% Tween-20, washed 3 times in TBS buffer with 0.1% Tween-20 for 2 minutes each, and incubated with primary antibodies at 4 °C overnight. The following day, membranes were washed 3 times with TBS with 0.1% Tween-20 for 3 minutes each and incubated with secondary antibody in 5% skim milk in TBS with 0.1% Tween-20 for 45 minutes at room temperature. Membranes were washed 5 times with 0.1% Tween-20 5 minutes each then imaged using ChemiDoc Imaging system (BIO-RAD).

### Immunofluorescence

Cells were grown on coverslips, washed twice with 1X PBS and fixed with 4% paraformaldehyde for 10 minutes at 37°C. The cells were quenched using 0.1 M Glycine in 1XPBS for 10 minutes, permeabilized using 0.1% Triton X-100 in PBS for 5 min and blocked with 1% fetal bovine serum (FBS), 1% bovine serum albumin (BSA) in PBS for 45 minutes. Primary staining was conducted in the blocking solution for one hour at room temperature, washed with 1XPBS-Tween 3 times for 5 minutes, and incubated with secondary in blocking solution for 1 hour. The coverslips were rinsed 3 times for 10 minutes each with 1XPBS-Tween and once with Millipore water then mounted using Anti-Fade (Invitrogen).

### Quantification of ER tubulation

Random ROIs were chosen per cell and subjected to analysis to differentiate between ER sheets and tubules. Diagonal lines were drawn per ROI, and each ER tubule intersecting the line was quantified as such based on three criteria: 1) the amplitude of each peak to baseline that is greater than or equal to a gray value of 50; 2) the peak baseline must reach a gray value of at least 10 on either side; and 3) the peak cannot be multiplet.

### Cell Culture and Transfection

COS-7, MIA PaCa-2 (MP2), Panc1 WT, Panc1 S237A, Panc1 shRNA control, Panc1 shTAK1, MEF WT and MEF αTAT1 KO were cultured in DMEM (GIBCO) supplemented with 10% fetal bovine serum (FBS) (GIBCO). Panc1 lenticontrol and Panc1 shTAK1 stables were also supplemented with puromycin (10ug/mL). Cells were kept in T-75 culture flasks in a 37 °C incubator with 5% CO_2_.

### Immunoprecipitation Assays

Cells were washed with 1XPBS and lysed with lysis buffer (20 mM HEPES [pH 7.4], 150 mM NaCl, 2 mM EDTA, 10 mM NaF, 10% [w/v] glycerol, 1% NP-40) for 20 minutes on ice. Cell lysates were transferred into microcentrifuge tubes and centrifuged at 13,000 RPM for 15 minutes. Supernatants were collected and incubated with appropriate antibodies and protein A agarose or protein B agarose for 4-6 hours or overnight at 4°C. Immunoprecipitated beads were washed three times with lysis buffer and stored in 2X sample buffer prior to western blot analyses.

### Crystal Violet Proliferation Assay

PANC-1 cells were plated at in 12-well plates. Cells were fixed at indicated time points (4% paraformaldehyde in PBS for 12 min). Following fixation, cells were washed, then stained with 0.1% crystal violet for 20 min. Cells were washed 3X, then air dried for 30 min prior to destaining (10% acetic acid, 90% water) for 20 min, and absorbance readings taken at 590 nm in a microplate reader.

### In gel digestion

To determine changes in the PANC-1 proteome upon αTAT1 S237A mutation, cell lysates were resolved on a 10% SDS-PAGE gel and stained with Bio-Safe Coomassie G-250 Stain. Each lane of the SDS-PAGE gel was cut into five slices. The gel slices were subjected to trypsin digestion and the resulting peptides were purified by C18-based desalting exactly as previously described^13,14^.

### Mass spectrometry and Data Search

HPLC-ESI-MS/MS was performed in positive ion mode on a Thermo Scientific Orbitrap Fusion Lumos tribrid mass spectrometer fitted with an EASY-Spray Source (Thermo Scientific, San Jose, CA). NanoLC was performed exactly as previously described (^13,14^). Tandem mass spectra were extracted from Xcalibur ‘RAW’ files and charge states were assigned using the ProteoWizard 3.0 msConvert script using the default parameters. The fragment mass spectra were searched against the mus musculus SwissProt_2018_01 database (16965 entries) using Mascot (Matrix Science, London, UK; version 2.6.0) using the default probability cut-off score. The search variables that were used were: 10 ppm mass tolerance for precursor ion masses and 0.5 Da for product ion masses; digestion with trypsin; a maximum of two missed tryptic cleavages; variable modifications of oxidation of methionine and phosphorylation of serine, threonine, and tyrosine. Cross-correlation of Mascot search results with X! Tandem was accomplished with Scaffold (version Scaffold_4.8.7; Proteome Software, Portland, OR, USA). Probability assessment of peptide assignments and protein identifications were made using Scaffold. Only peptides with ≥ 95% probability were considered. Label-free peptide/protein quantification and identification. Progenesis QI for proteomics software (version 2.4, Nonlinear Dynamics Ltd., Newcastle upon Tyne, UK) was used to perform ion-intensity based label-free quantification as previously described.

### CRISPR genome editing and knock-in

Parental Panc1 cells were first transfected using RNAiMAX reagent (Thermo DNA Technologies) with ribonucleoprotein particles (RNPs) consisting of a crRNA (guide RNA CTTACATTCTCGGTCACTAG), universal 67mer tracrRNA, and Cas9 protein (Integrated DNA Technologies). To estimate cutting efficiency, we quantitated the proportion of abnormal DNA fragments generated by the procedure using DNA prepared from the transfected cell population and a polymerase chain reaction (PCR)-based T7 endonuclease assay (New England BioLabs). The PCR was performed with primers flanking the predicted ligation-junction product (forward primer AGAATACCAGGGGGCCATGA and reverse primer AACGAATCCTTTCCCTGGGTC). We then co-transfected Panc1 cells with CRISPR RNPs and single stranded oligodeoxynucleotide repair template (CCAGCTCCAGCAAGGAAGCTGCCACCCAAGAGAGCAGAGGGAGACATCAAGCCAT ACTCC**GCA**AGTGACCGAGAATGTAAGAGGGGCAAGGGTCGGGTGTCTGGGCCTGGGGTACCTTAACAC; the site for the serine to alanine amino acid substitution is shown in bold typeface), which would substitute alanine (GCA) for the serine amino acid (TCT) upon microhomology-based DNA end-joining. After two days, single cells were deposited in 96-well plates by flow sorting with a FACS Aria III (BD Biosciences). Single clones were expanded and DNAs were prepared from 81 clones. Clones carrying GCA mutations were identified by PCR and DNA sequencing with the reverse primer. No homozygotes were identified on the first trial; consequently, we chose a clone that was heterozygous TCT/GCA and repeated the procedure and screened another 96 clones and identified two clones that were homozygous TCA/TCA by PCR-sequence analysis.

### Statistical Analysis

Statistical analysis was performed using an unpaired two-tailed student t test. Significant statistical differences between groups were indicated as: *P < 0.05. Data are presented as mean ± SEM. No statistical method was used to predetermine sample size. No data was excluded from the analyses. Statistical analyses and graphics were carried out with GraphPad Prism software and Microsoft Excel.

## Acknowledgements

We would like to thank the University of Arizona Cancer Center for their assistance. This work was supported by NIH grant R35GM148171 awarded to N.Y.L, and the Cancer Center Support Grant (P30 CA023074).

## Author Contributions

H.R.O., P.C.F., A.R., and T.A. conducted majority of the experiments and data analysis; H.R.O. and P.C.F. wrote the manuscript; N.A.E. generated the CRISPR cell lines; P.R.L. and K.M. assisted with proteomics analysis; N.Y.L. conceived the study, analyzed all data, prepared, and edited the manuscript.

## SUPPLEMENTARY FIGURES

**S1.**
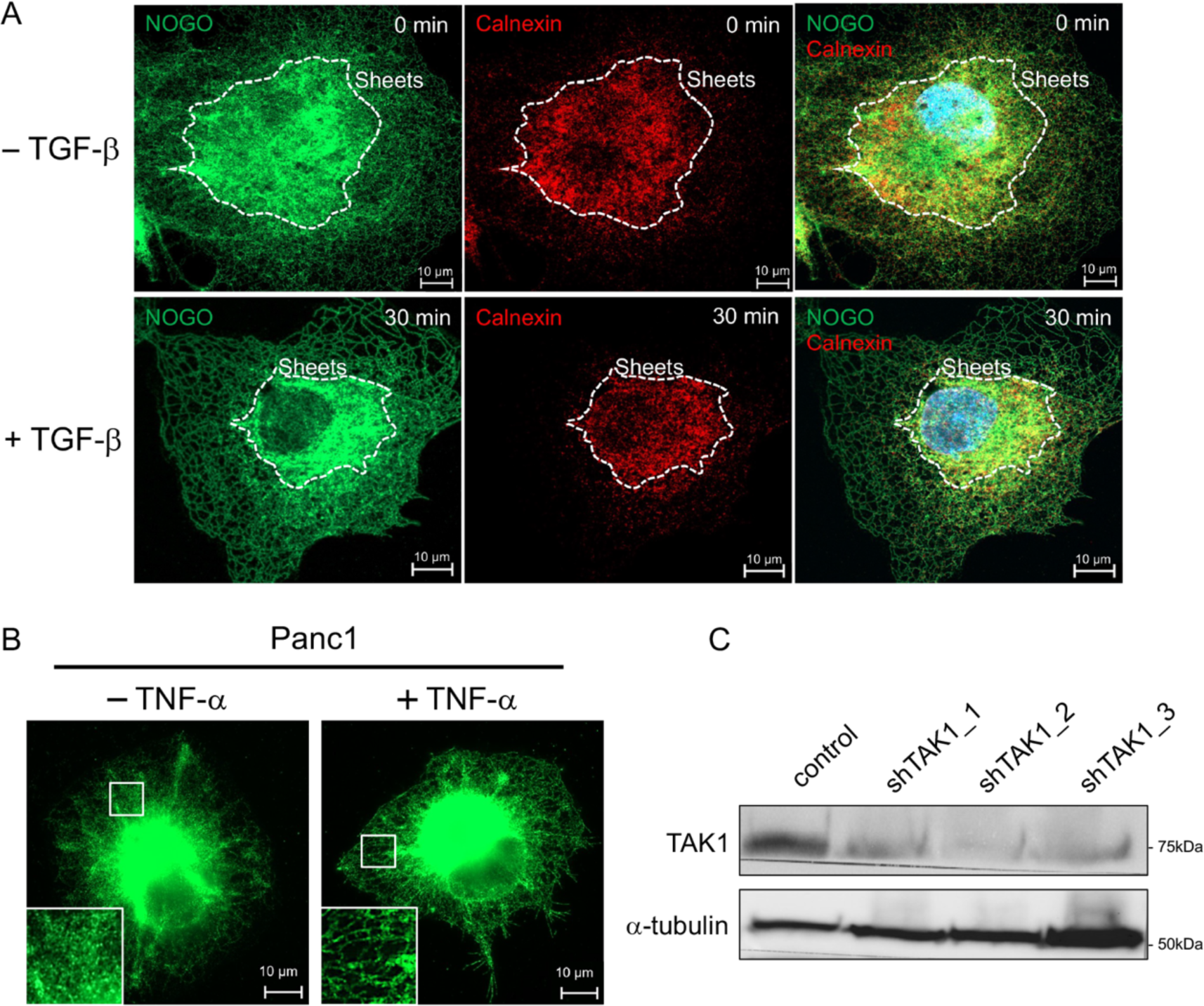
A) Representative immunofluorescence images of COS-7 cells dual-stained with NOGO (green) and calnexin (red) upon treatment with or without TGF-β (200 pM) for 30 min. White dotted lines depict the boundaries of calnexin distribution. B) Representative images of immunofluorescence staining of NOGO (green) in Panc1 cells, subjected to TNF-α (20 ng/mL) stimulation for 30 min prior to staining. C) Western blot shows endogenous TAK1 depletion using three different shRNA against human TAK1 in PANC-1 cells showing total α-tubulin as a control.

**S2.**
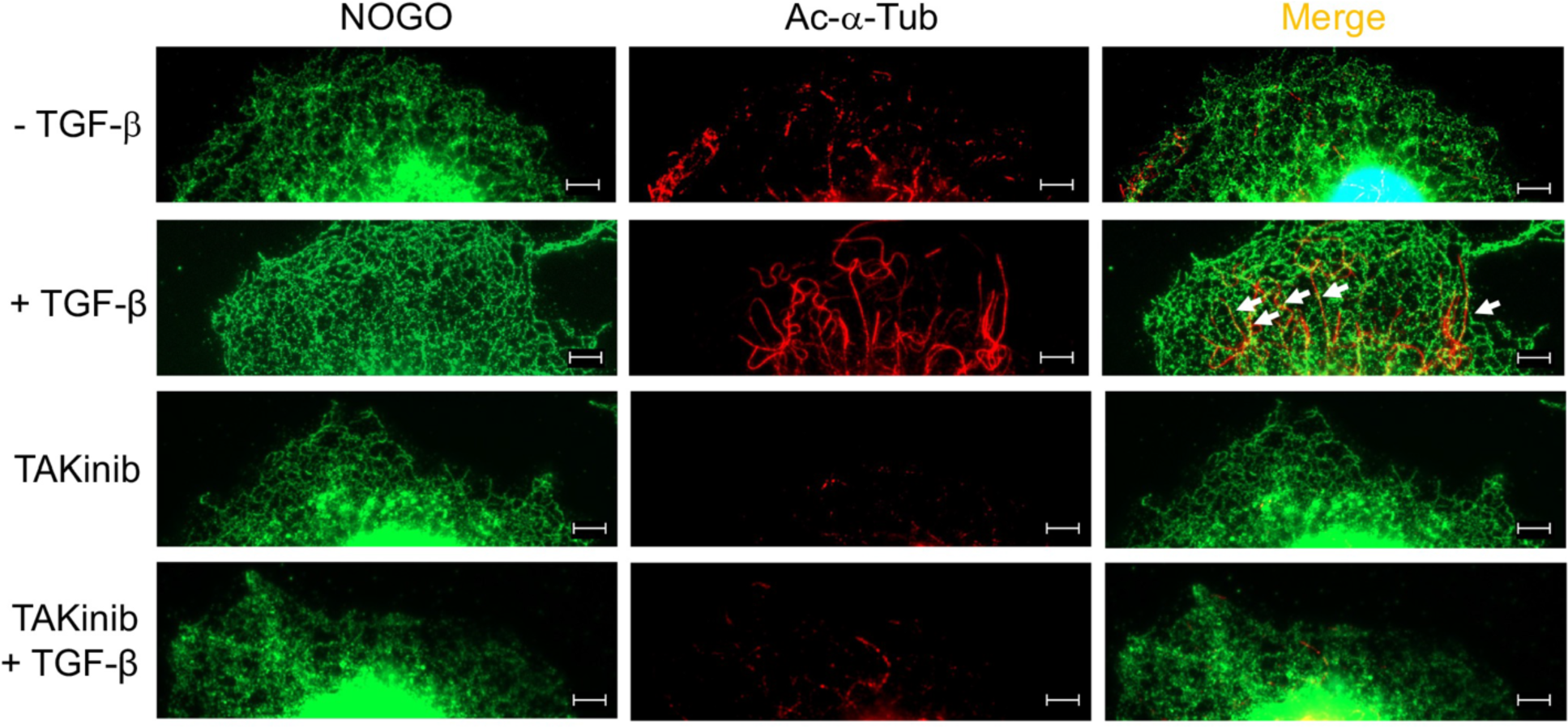
Representative images of immunofluorescence staining of NOGO (green) and acetyl-α-tubulin (Red) in Panc1 cells, subjected to pretreatment using a pharmacological inhibitor TAKinib (10 µM) for 30 min then addition of TGF-β (200 pM) for 30 min. ER sliding along acetylated MT is indicated by white arrows.

**S3.**
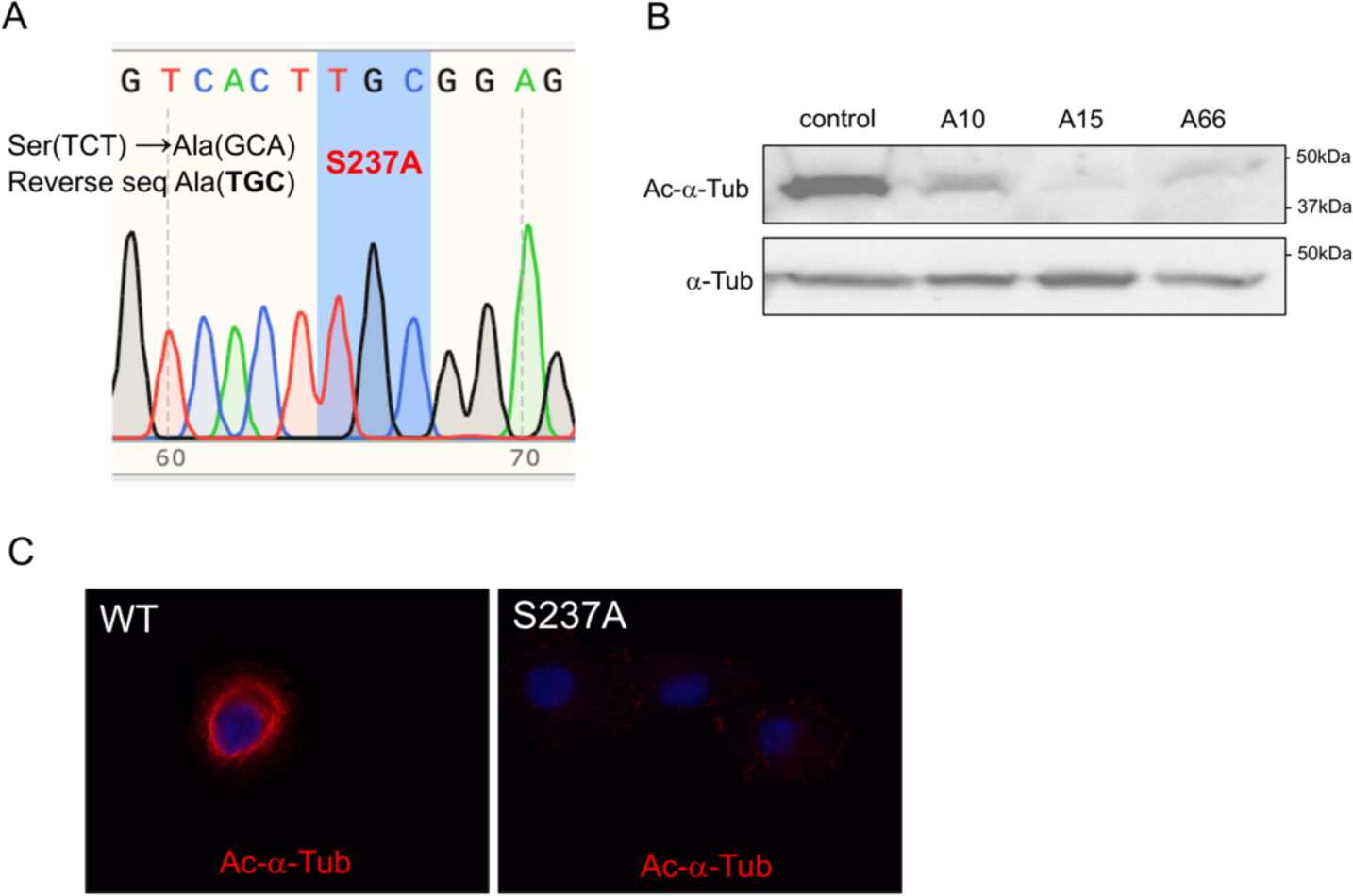
A) Sequence chromatogram depicts the nucleotide change from Ser to Ala in the Panc1 αTAT1 S237A cells. B) Western blot shows endogenous depletion of acetyl-alpha-tubulin in three different CRISPR knock in clones in Panc1 cells. C) Representative images of immunofluorescence staining of acetyl-alpha-tubulin (red) in Panc1 WT and αTAT1 S237A cells. Clones A15 and A66 were used interchangeably throughout the studies.

**S4.**
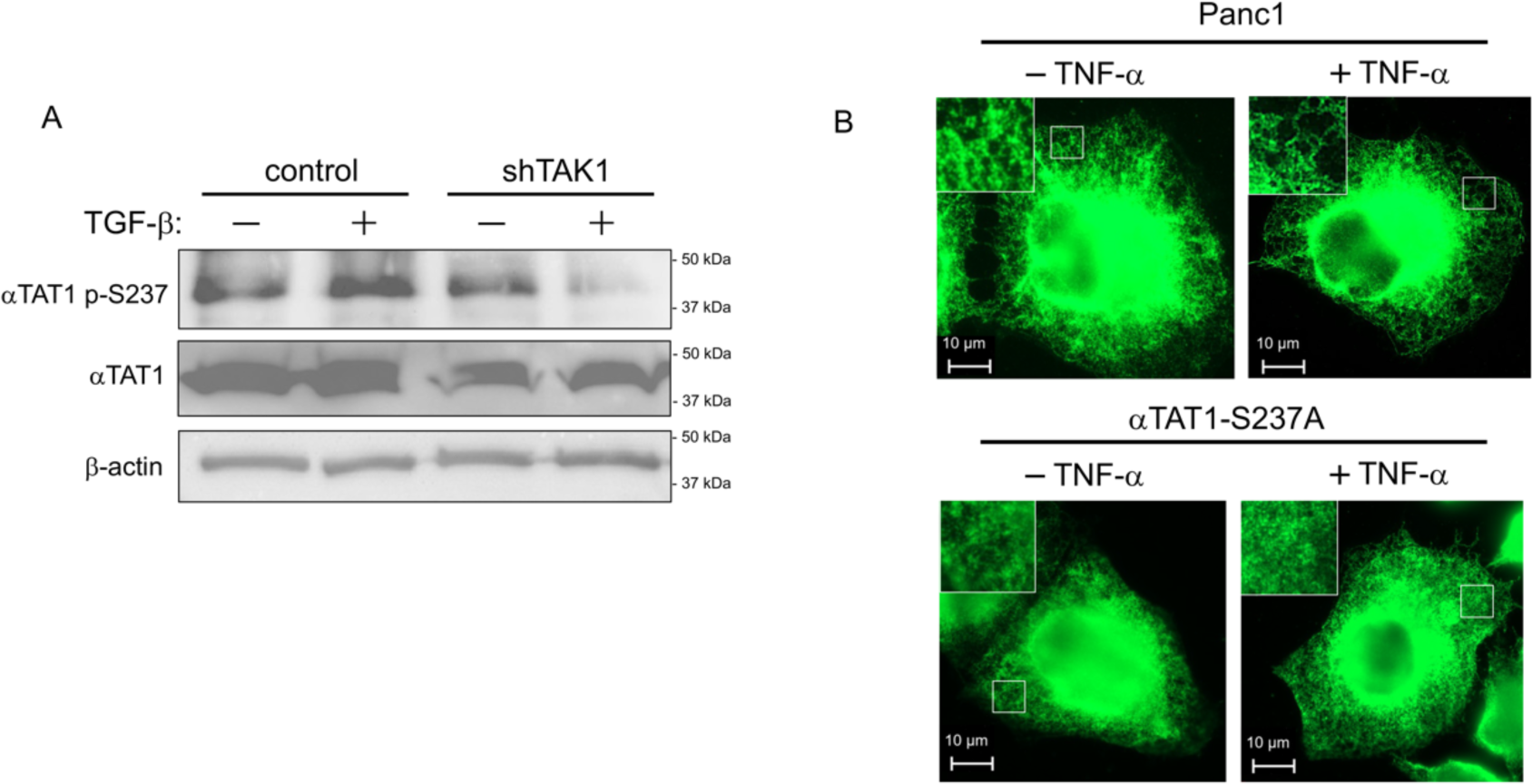
A) Western blots for Panc1 WT and shTAK1 cells subjected to 30 min TGF-β (200 pM) showing αTAT1 S237 phosphorylation using a phosphoantibody as well as levels of total αTAT1 and β-actin. B) Representative images of immunofluorescence staining of NOGO in Panc1 WT and Panc1 αTAT1 S237A cells, treated with and without TNF-α (20 ng/mL) for 30 min prior to staining.

**S5.** MS-Quantitative proteomics raw data showing the statistically significant responders when comparing Panc1 WT versus αTAT1 S237A.

**S6.**
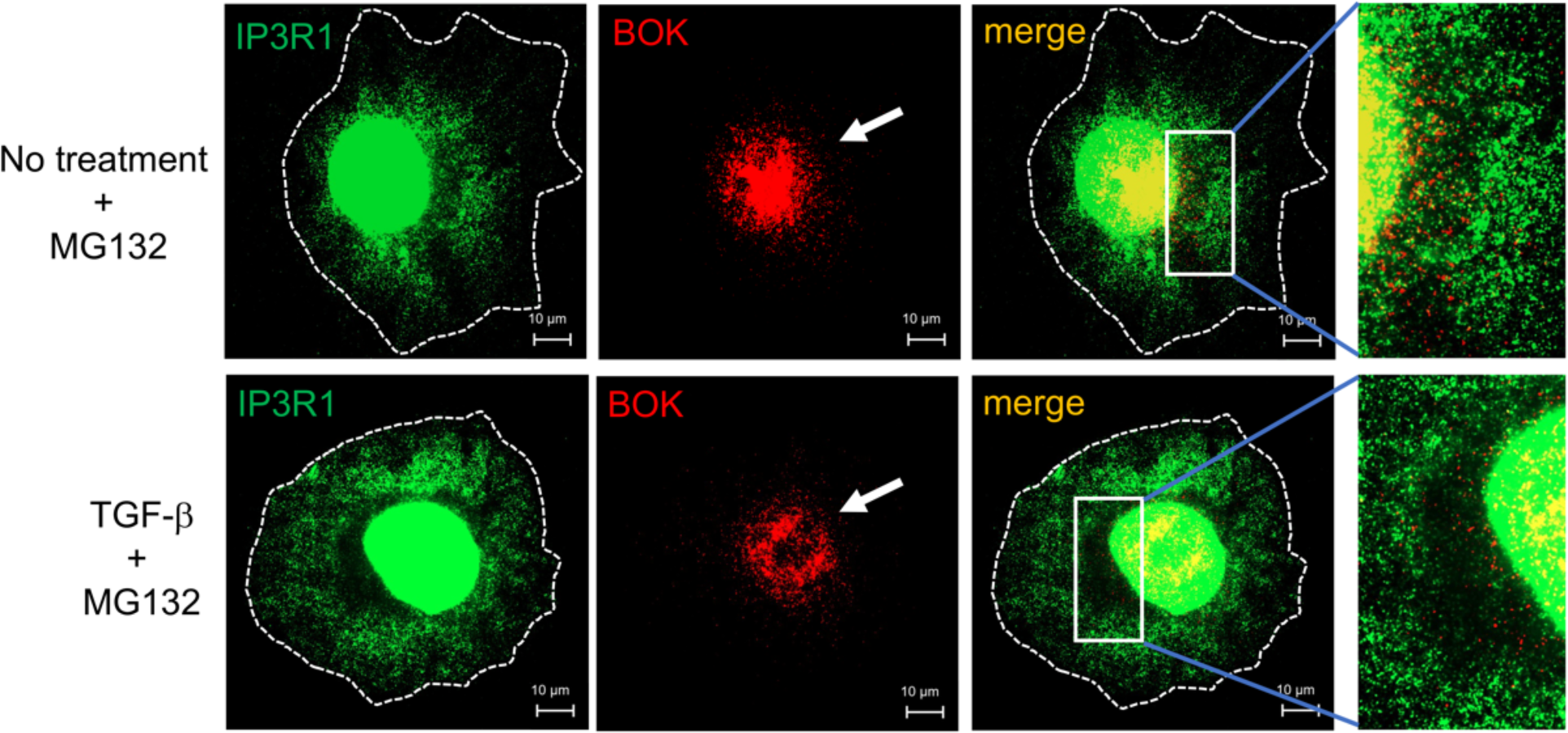
Representative images of immunofluorescence staining of IP3R (green) and BOK (Red) in Panc1 WT previously pretreated with proteasome inhibitor MG132 (25 µM) for 30 min as well as TGF-β stimulation (200pM) for 30 min. BOK cellular localization is signaled by white arrows.

**S7.**
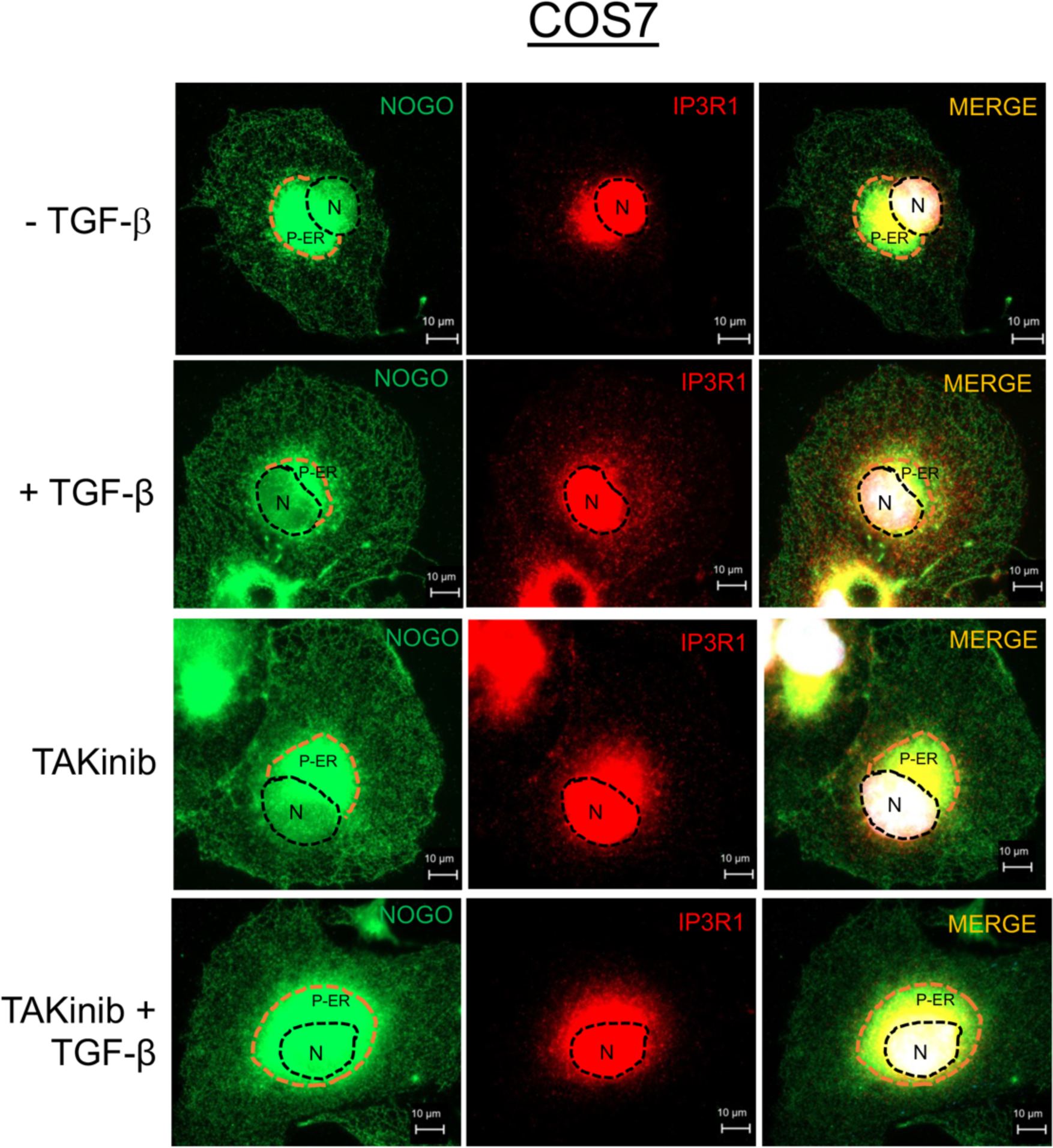
Representative images of immunofluorescence staining of NOGO (green) and IP3R (Red) in COS-7 cells pretreated with TAKinib (10 µM) for 30 min as well as with TGF-β stimulation (200 pM) for 30 min prior to staining.

**S8.**
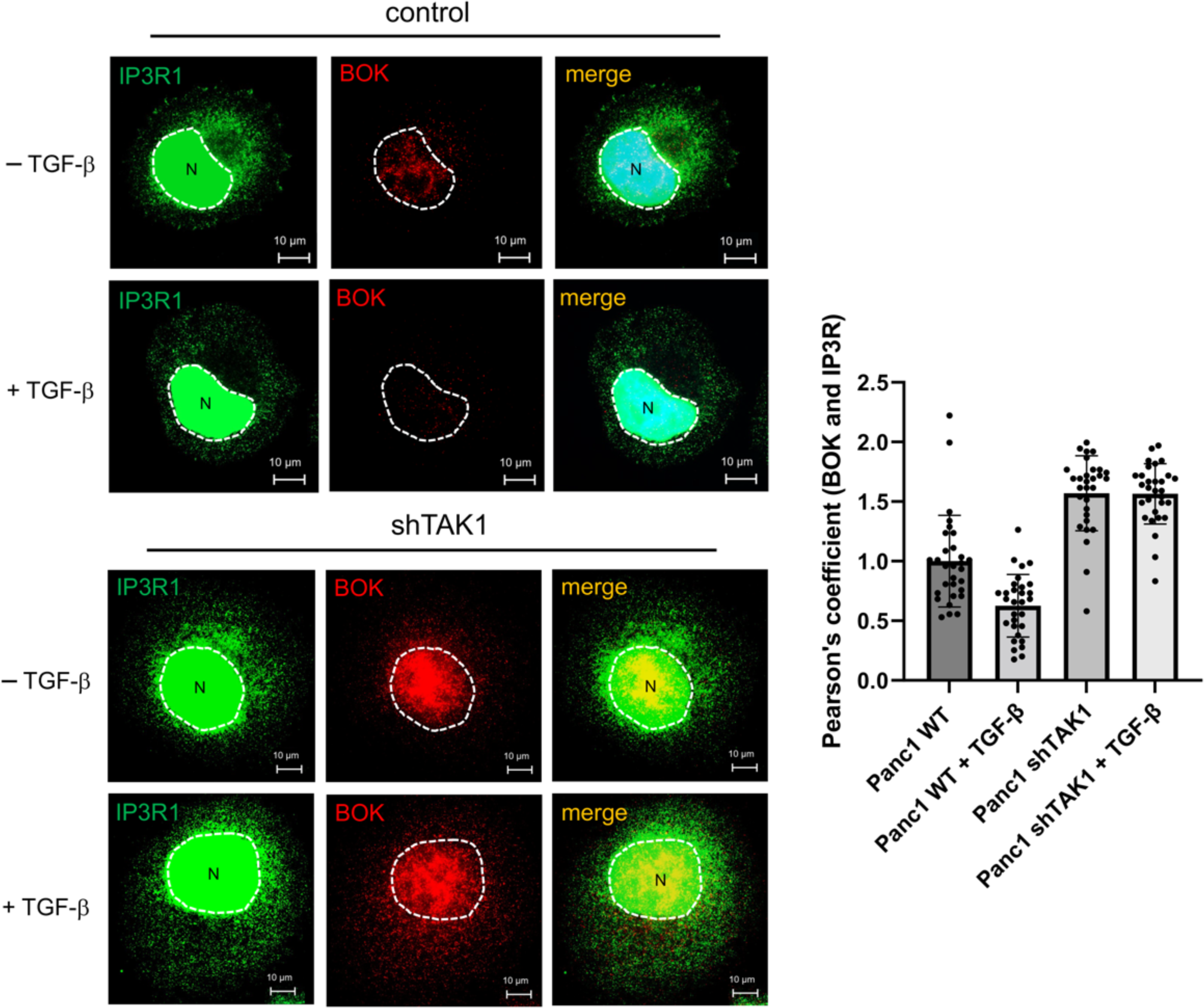
Representative images of immunofluorescence staining of IP3R (green) and BOK (Red) in Panc1 control and shTAK1 cells treated with and without TGF-β (200pM) for 30 min.

